# CYCLeR– a novel tool for the full isoform assembly and quantification of circRNAs

**DOI:** 10.1101/2021.04.27.441578

**Authors:** Stefan R. Stefanov, Irmtraud M. Meyer

**Affiliations:** Berlin Institute for Medical Systems Biology, Max Delbrück Center for Molecular Medicine in the Helmholtz Association, Hannoversche Str. 28, 10115 Berlin, Germany; Freie Universität Berlin, Department of Biology, Chemistry and Pharmacy, Institute of Chemistry and Biochemistry, Thielallee 63, 14195 Berlin, Germany

## Abstract

Splicing is one key mechanism determining the state of any eukaryotic cell. Apart from linear splice variants, circular splice variants (*circRNA*s) can arise via non-canonical splicing involving a *back-splice junction* (BSJ). Most existing methods only identify *circRNA*s via the corresponding BSJ, but do not aim to estimate their full sequence identity or to identify different, alternatively spliced circular isoforms arising from the same BSJ. We here present CYCLeR, the first computational method for identifying the full sequence identify of new and alternatively spliced *circRNA*s and their abundances while simultaneously co-estimating the abundances of known linear splicing isoforms. We show that CYCLeR significantly out-performs existing methods in terms of sensitivity, precision and quantification of transcripts. When analysing *D. melanogaster* data, CYCLeR uncovers biological patterns of circRNA expression that other methods fail to observe.

## 1 Introduction

One major source of complexity in eukaryotes is splicing whereby one gene can give rise to a number of splicing products depending on the cell’s state (tissue, developmental stage, disease state etc). Splicing not only gives rise to linear splicing isoforms, but — interestingly — has also been recently shown to yield so-called circular splice variants or isoforms (*circRNA*s). These arise via a so-called *back-splice junction* when a downstream 5’ splice site is covalently linked to an upstream 3’ splice site (6). The corresponding junction is called a back-splice junction (BSJ).

For decades, scientists have focused almost exclusively on linear splice variants and their functional roles. This focus, however, has recently shifted towards *circRNA*s (7; 8; 9; 10; 11). While the molecular functions of select *circRNA*s have already been identified (8; 12; 13; 14; 7), we are only beginning to understand the mechanisms by which circRNAs arise and the functional roles they play *in vivo*.

Since *circRNA*s constitute a form of alternative splicing of the nascent RNA transcript (7), the raw RNA-seq reads deriving from linear and circular splicing isoforms of one and the same gene cannot be easily used to reliably infer any new *circRNA*s and their abundance, see Figure 1. Only RNA-seq reads spanning a *back-splice junction* provide direct evidence for *circRNA*s. These reads, however, provide only limited information on the full sequence identity of the underlying *circRNA*s, thus making the proper identification and quantification of new *circRNA*s impossible.

**Figure 1:**
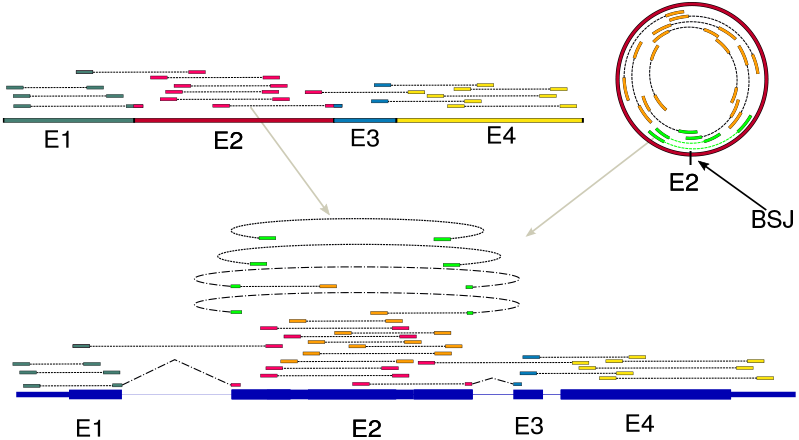
Challenge of identifying *circRNA*s from RNA-seq data. Typical, raw transcriptome data from linear and circular splicing isoforms (top left and right) comprises a multitude of pair-end reads covering the exons of these isoforms (E1 etc, colouring of pair-ed reads according to the exon from which they derive). In order to infer the original splicing products from these raw transcriptome reads, they are typically first mapped to the genome (bottom). Most of the mapped reads will not cover splice sites (exon-intron boundaries) and could either derive from a linear and circular splicing isoform. One challenge is that only reads spanning a *back-splice junction* provide direct evidence for *circRNA*s (marked in light green). As is also clear from this picture, the correct identification and quantification of *circRNA*s cannot be achieved without the simultaneous identification and quantification of the linear splicing isoforms. Thus, if the linear splicing isoforms of a gene are known up-front, their correct quantification needs to be estimated in conjunction with the identification and correct quantification of unknown *circRNA*s.

The ultimate goal in transcriptome assembly is to identify the full sequence identify of all linear and circular splice variants and to estimate their relative abundances. The relative expression levels are key requirements for differential expression analyses and co-expression correlation analyses. Most existing methods for transcriptome assembly, however, ignore *circRNA*s and only focus on linear splicing isoforms. Commonly used transcriptome assembly programs (15; 16) are based on creating a directed acyclic splice graph, whose nodes and edges are determined by the presence of forward junction spanning reads in the raw transcriptome data. These programs, however, cannot handle the cyclic splice graphs that the *circRNA*s represent.

One added challenge is that *circRNA*s typically constitute only a small fraction of the transcriptome (1-3% of linear poly-A transcripts in common cell lines (17)), thus making the identification of *circRNA*s via an increased RNA-seq library depth no viable option. In addition, the expression of linear splicing isoforms from the same gene locus can significantly skew the circular transcript assembly. For efficient *circRNA* detection, there is thus the need for an enrichment of *circRNA*s in a sample. Due to the lack of free 5’- and 3’-ends, *circRNA*s are resistant to exonuclease enzymes (18; 7). By using an exonuclease treatment, transcriptome libraries can thus be enriched in *circRNA*s. As alternative, *circRNA*-enriched libraries can also be produced by poly-A depletion (19). These circRNA-enriched libraries, however, help circRNA-identification, but cannot be used to improve circRNA-quantification as the depletion steps are known to affect different isoforms differently (2). A combination of treatments can lead to an almost exclusive circular RNA-seq library(5).

Alternatively, *circRNA* isoforms can be discovered by nanopore-sequencing linear molecules that were generated via a rolling circle amplification of the corresponding *circRNA* template. Also this procedure first requires linear transcript depletion ahead of the rolling circle amplification. This strategy for *circRNA* full-isoform identification, however, has issues with reproducibility and requires a high number of replicates to identify all *circRNA* isoforms (1).

In order to enable a fair assessment of CYCLeR compared to existing tools for *circRNA* identification and quantification, it is key to first classify the existing methods based on their goals. We define class I to comprise well-known *circRNA* tools that aim to identify and quantify *circRNA*s solely on reads spanning BSJs. Unlike CYCLeR, these tools do not aim to identify the full-sequence identity of new *circRNA*s, let alone multiple, alternatively splicing *circRNA*s corresponding to the same BSJ (17).

Class II contains *circRNA* characterisation tools that take as input the predictions generated by methods from class I in order to produce a set of potential splicing events for each sequence interval defined by a BSJ. Since the alternative splicing of linear RNAs may obstruct the detection of circular alternative splicing (AS) events, class II tools require *circRNA* enriched libraries to function properly. Two approaches to detect the alternative splicing of *circRNA*s from transcriptome RNA-seq data have emerged. The first approach is conceptionally based on exon and alternative splicing detection, using mate-pair information of pair-end reads, spanning a BSJ (20; 21). This approach, however, can only detect AS events within the range of the insert size used for making the RNA-seq library. The second approach compares circle enriched and control libraries, similarly to CYCLeR. This strategy has the advantage of removing the dependency on the library insert size and is employed in the CIRCexplorer2 (19) pipeline. Reads spanning a BSJ typically represent only around 0.1% of a library, thereby making quantification based solely on those reads unreliable.

This challenge prompted the release of special quantification tools (defined as class III) that take as input the output of class II or class I tools and produce as output estimated circRNA levels. The tools of class III can be divided into two sub-classes. One sub-class (class IIIa) comprising tools that provide BSJ quantification as well as expression level ratios of the BSJ and FSJ in the locus of the same *circRNA* (25; 2). While those values are reportedly in agreement with qPCR results (25; 2), they do not allow to deduce the relative expression levels of all alternatively-spliced linear and/or circular isoforms that happen to overlap the same BSJ and FSJ, respectively, which is what our method CYCLeR is aiming to do. An interesting case is sailfish-cir (23), the sole member of the sub-class IIIb. sailfish-cir takes as an input a list of BSJs and known linear transcripts and makes a pseudo-linear reference of potential *circRNA* transcripts. sailfish-cir later uses a combination of the pseudo-linear reference and known linear transcripts to quantify linear and circular transcripts simultaneously. To conclude, all class III tools provide a circRNA abundance estimation that is only based on a BSJ, but ignore any additional, alternatively spliced *circRNA* isoforms that may be sharing the same BSJ.

The tools (22; 24) which constitute class IV aim to recover the full-sequence identity of *circRNAs*. One example is CIRI-FULL which employs an extension of the mate-pair approach of the class II tools in order to derive quantitative information on full-length *circRNA* isoforms. The recommended input to CIRI-FULL are 2 x 250 nt paired-end libraries which allows for the full sequence-identity recovery of some *circRNA*s. This insert size used for making the RNA-seq library thus limits the scope of the algorithm’s output, thereby allowing only for the identification of circular isoforms of up to 600 nt length (22). The assembly strategy applied in CircAST is less ambitious as the tool does not utilise quantitative information for the reconstruction (24). Class IV tools provide relative abundances of *circRNA* isoforms as counts based on fractions of the BSJ spanning reads. All class IV tools require *circRNA* enriched libraries as input.

To conclude, the existing methods for *circRNA* investigation constitute a wide set of diverse tools with distinct goals and features, see classes I to IV defined above and Table 1 and Table 2. Right now, however, there exists no method for identifying the full-sequence identity of *circRNA*s as well as their potential alternatively spliced variants and for simultaneously estimating the expression levels of known linear splicing isoforms. This is why were are introducing here our method CYCLeR (Co-estimate Your Circular and Linear RNAs).

**Table 1:**
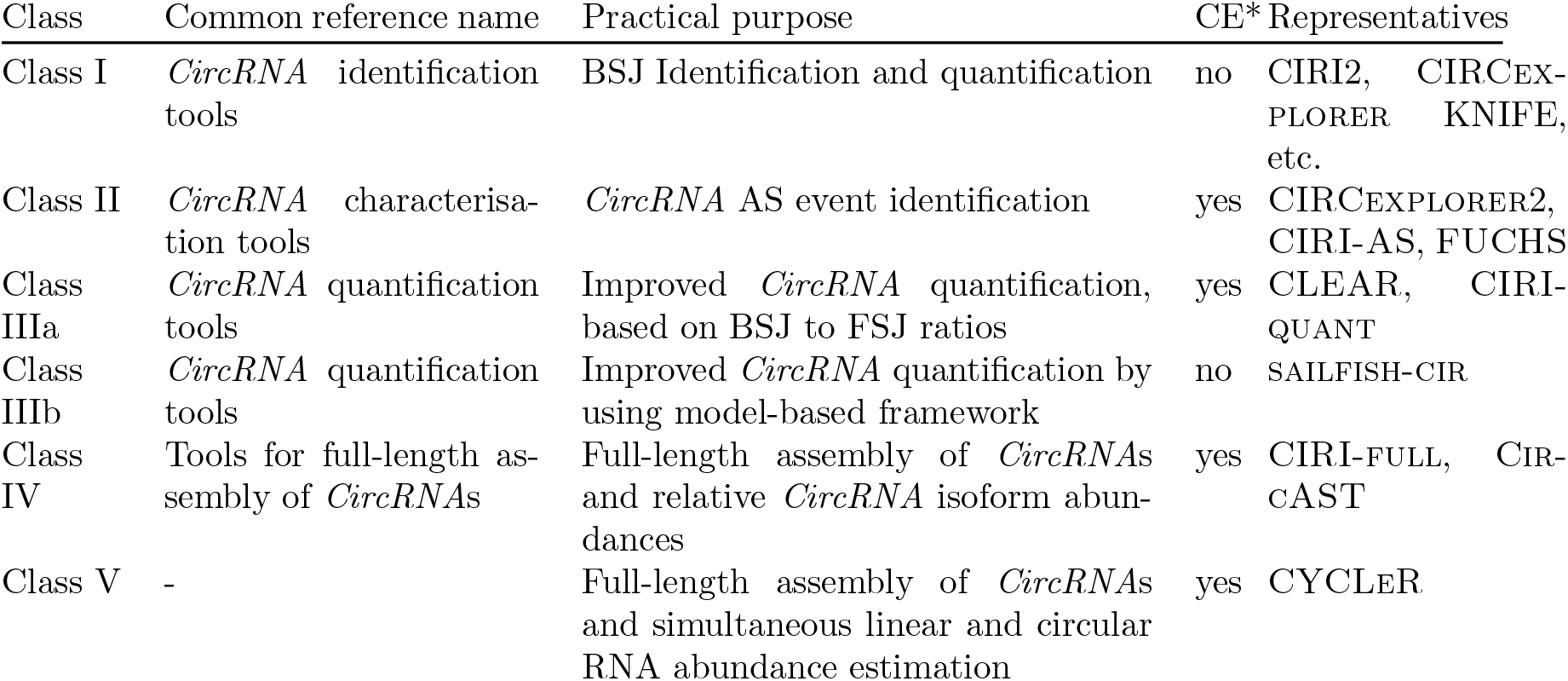
**Classification of existing methods for *circRNA* identification and quantification** according to their goals and the input they require. Tools in column CE (Circle Enriched) denoted by a ‘yes’ require *circRNA* enriched libraries for optimal performance.

**Table 2:**
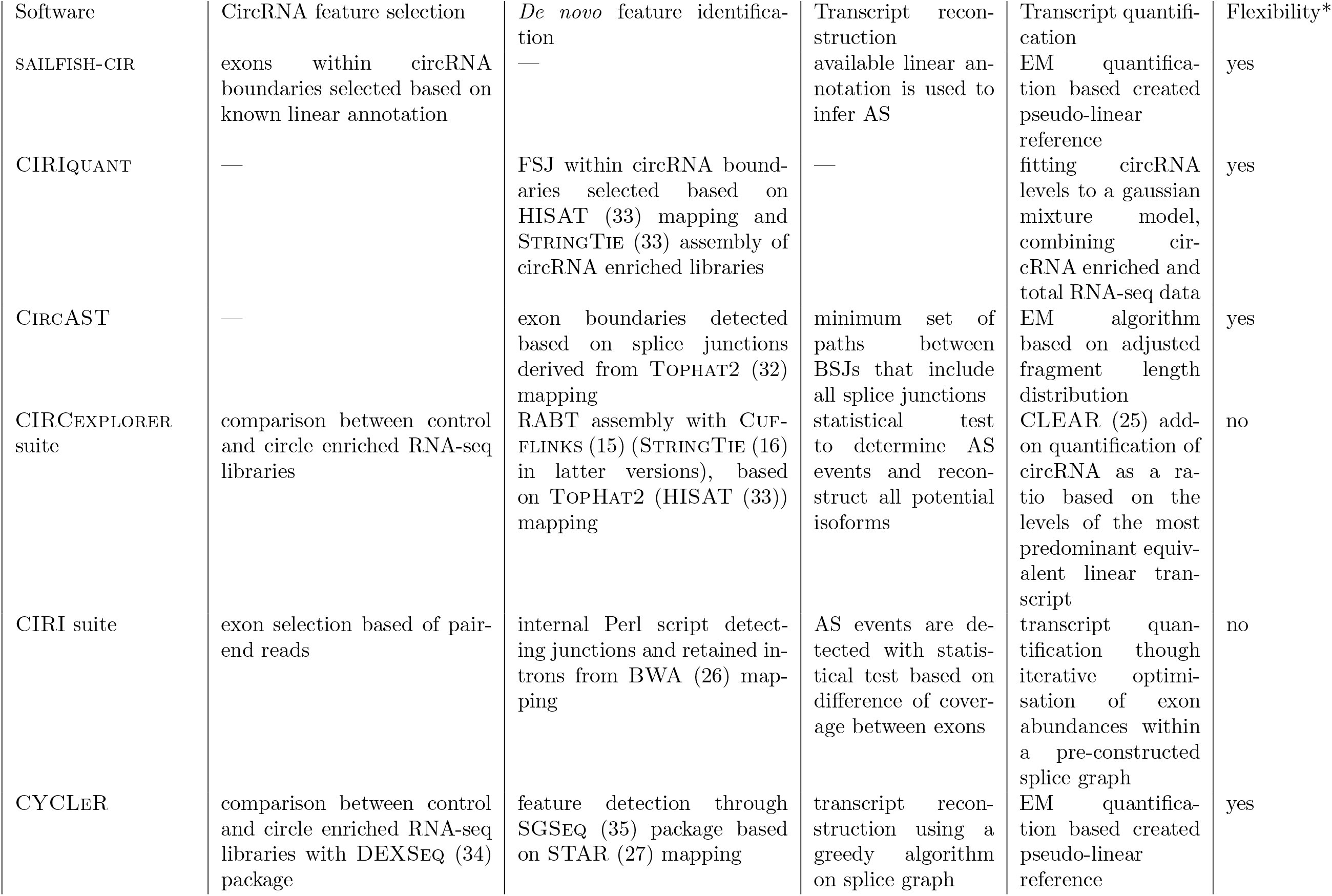
Overview of relevant circRNA transcript reconstruction and quantification tools. Abbreviations used: AS - alternative splicing, BSJ *-back-splice junction*, EM - expectation maximisation; * indicates that the tool is fully compatible with various BSJ identification tools

By comparing default (control) and circle-enriched RNA-seq libraries as input, the algorithm underlying CYCLeR first captures circle-specific features and then reconstructs full-length *circRNA* isoforms via a flow-based algorithm. The *circRNA* transcripts are then converted into a pseudo-linear isoform profile in order to estimate the linear and circular transcripts’ abundance via expectation-maximisation.

As our method has several unique features, we benchmark different aspects of it by comparing it to the most adequate state-of-the-art methods that share the same feature. We thus assess the assembly of *circRNA*s in CYCLeR separately from isoform quantification. For the assembly benchmarking, we compare CYCLeR against class IV tools as well as CIRCexplorer2 from class II. This is because CIRCexplorer2’s isoform reconstruction module deduces all potential isoforms involving the identified *alternative splicing* events (and thus does not account for correlated pairs of *alternative splicing* events such as the skipping of two neighbouring exons, or mutually exclusive exons). For benchmarking the isoform quantification of CYCLeR, only class IV tools are considered. Note that class III cannot be utilised for this purpose as they have no ability to distinguish multiple circular isoforms sharing the same BSJ. For an overview of relevant existing methods, please refer to Table 1 and Table 2.

## 2 MATERIALS AND METHODS

In this section, we introduce the steps and algorithms employed by CYCLeR. CYCLeR takes as an input a set of BSJs, control and circRNA enriched libraries and – optionally – annotation information on linear transcripts. CYCLeR first prepares gene specific splice graphs and iteratively reconstructs potential isoforms. After CYCLeR assembles the isoforms from every sample, it prepares a complete combined set of isoforms to serve as a reference for quantification.

We also explain the reasoning behind the parameters of the different benchmarking strategies. We test assembly efficacy as well as quantification accuracy, as is common, with a simulated dataset. We also analyse real transcriptome data to showcase the difference of the output produced by class III tools and by CYCLeR and to highlight the advantage that full-sequence information brings to an analysis. A qPCR benchmark was used to show the advantage of CYCLeR with respect to class IIIa tools. We also conducted a exploratory transcriptome analysis to highlight the advantages of CYCLeR over class IIIb tools.

### 2.1 Selection of a reliable BSJ set

The detection of BSJs relies on the capability of a mapper to detect chimeric reads. Different mappers detect different sets of chimeric reads per sample. To be certain that a reliable set of BSJ sites has been identified, we use CIRI2 and CIRCexplorer2 as input. They employ BWA-MEM and STAR for chimeric detection, respectively. We also allow the option of a BSJ set to be added manually as an input via a TSV file. Since CYCLeR requires circRNA enriched libraries as input, we can easily adjust for false positives by comparing the BSJ set identified from total RNA-seq to the set identified from circRNA enriched RNA-seq.

### 2.2 Creation of circular splice graphs

CYCLeR utilises the two-pass mapping mode of STAR (27) to recover all split-mapped reads. These mapped reads are then used to identify both annotated and new genomic features such as exons, retained introns and splice junctions. For each gene, CYCLeR subsequently creates a corresponding splice graph that also contains information on feature abundances (needed for the reconstruction algorithm). Then, only the genomic features that fall between a BSJ-start and a corresponding BSJ-end coordinate (*i*.e. a circle generating loci) get extracted and a circle-specific splice graph is constructed. The corresponding features (exons and splice-site junctions) within this graph are then re-quantified, normalised and adjusted for GC-content and length biases. Features that are depleted in *circRNA*-enriched samples are detected either through a direct quantity comparison or a negative binomial test and excluded from the final splice graph to minimise the number of false reconstructions later on, see Figure S3 in the supplementary information.

### 2.3 Reconstruction of circRNA transcripts

One common approach towards transcriptome assembly is to use a so-called flow-based algorithm to disentangle the splice graph (16).

In CYCLeR, we employ a greedy algorithm for the iterative reconstruction of transcripts to ensure a low number of false-positive-assembled sequences. To this end, we use the comprehensive splice graph, created in the previous step and start by selecting the exon with the lowest abundance. We then identify the maximum flow though this exon inside the splice graph and reconstruct the corresponding circular transcript. The corresponding exon abundances then need to be subtracted from the respective features of the original graph and any fully depleted features be removed. This operation is repeated until no more transcripts can be reconstructed, see Figure 2 for a detailed example.

**Figure 2:**
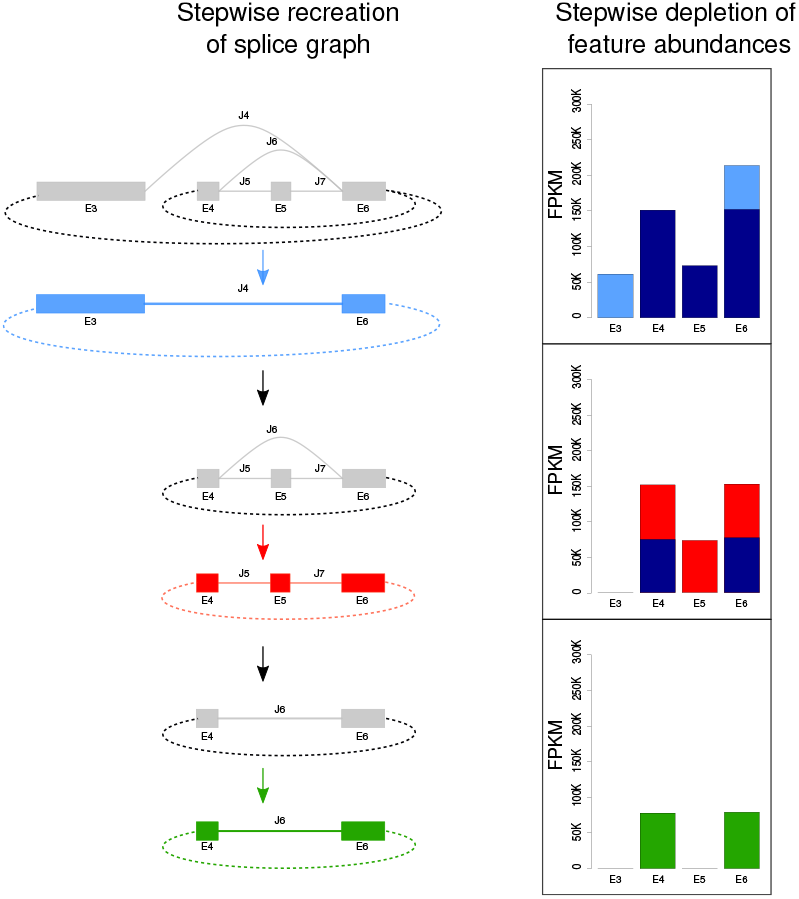
**Circle transcript reconstruction within** CYCLeR **for the example of the 5-HT2A gene.** Starting with the full splice graph for the entire gene locus and its respective exon abundances (see top line, left, in grey for the graph and the FPKM-plot at the top right), CYCLeR extracts the circle-specific sub-graph corresponding to splice-site junction J4 which falls between a BSJ-start and a corresponding BSJ-end coordinate (see second line from top left, in blue for the graph). This blue sub-graph correponds to a single circular splicing isoform. This blue subgraph and the corresponding exon abundances are subsequently subtracted from the original, full splice graph (see graph at the top left and middle FPKM-plot) to yield the remaining splice graph (third line from top, in grey). Similarly to before, CYCLeR then extracts the next circle-specific sub-graph, this time corresponding to a *back-splice junction*-spanning splice-site junction between exon E6 and E4 (fourth line from top, in red). This sub-graph provides evidence for a circular splicing isoform comprising three exons E4, E5 and E6 (note the different exon abundances). The quantities corresponding to this circular isoform are sub-sequently deleted from the remaining grey splice graph, resulting in a sub-graph (second line from bottom, left, grey) that corresponds to another circle-specific graph, this time comprising only exons E4 and E6, but not E5 (bottom line, in green). This sub-graph and its abundances provide evidence for a single circular isoform (bottom FPKM-plot).

### 2.4 Combining circRNA transcript sets across samples

Since different samples and replicates produce different sets of predicted transcripts, assembly tools often have modules for merging transcripts (15; 16). In CYCLeR, transcripts from different samples are merged into a single reference, while simultaneously using the differences between sets to fix discrepancies caused by mapping artefacts. The final set of transcripts can then be exported in terms of an annotation file (in GTF format or flat file) and a sequence file in FASTA format.

### 2.5 CircRNA transcript quantification

CYCLeR performs transcript quantification similarly to (12). For each *circRNA* transcript, a corresponding pseudo-linear reference is created, for details see Figure S5 and supplementary information section “CircRNA transcript quantification”. Afterwards, the set of pseudo-linear transcripts is combined with annotated linear transcripts and used to create an index for quantification with Kallisto (28).

### 2.6 Benchmarking with simulated data

The benchmarking of tools for transcriptome assembly and quantification is commonly based on dedicated, simulated data. Since CYCLeR aims to identify full-length circular isoforms and alternative splice variants and their abundance, we can only benchmark against the tools from class IV. In principle, CIRCexplorer 2 from class II can also be included in the benchmarking as it outputs potential transcripts derived from on all (!) potential combinations of AS events. The parameters used for the programs are summarised in Figure S12. Sensitivity and precision are calculated in the usual manner:

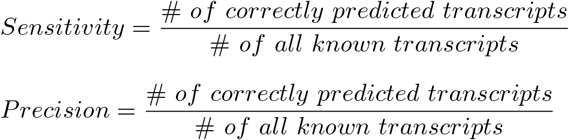

The quantification correlations are based on the estimated values of correctly assembled transcripts. The Pearson product correlation between estimated and simulated abundances is calculated differently with regard to the difference in the output. For CYCLeR, this is done as follows:

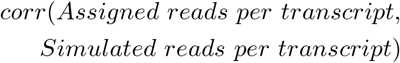

and CIRI-full as:

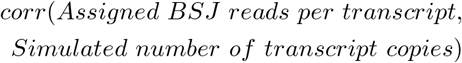

#### 2.6.1 Simulated dataset

The transcript reconstruction in CYCLeR requires as input a control and a *circRNA* enriched library, preferably as pair of replicates. CIRCexplorer 2 has similar requirements. The matepair-based methods require long library inserts with preferably 250 bases sequenced on both sides. With CYCLeR, we focus solely on library preparation involving RNA fragmentation in order to avoid a rolling circle amplification which could introduce unknown skewing. For the benchmarking of CYCLeR and existing methods, we thus simulate two types of RNA-seq libraries: one library with a median fragment length of a 230 bp and 75 bp sequencing and one library with a median fragment length of 480 bp and 250 bp sequencing. From both, we simulate a pair of control and circle-enriched libraries with two replicates each, for details see supplementary information section “Simulation of RNA-seq data” as well as Tables S1, S2 and S3 and Figures S1 and S2.

### 2.7 Benchmarking with real data

#### 2.7.1 Benchmarking with qPCR

A quantitative benchmarking involving qPCR with primers converging on one BSJ only makes sense for tools of class I and class IIIa which have no ability to detect alternative splicing isoform mapping to the same BSJ. Those tools output the relative abundance of a *circRNA* based on BSJ spanning reads and do not account of AS events that may occur within the locus of the same circle. Since CYCLeR does have the ability to quantify multiple, alternatively spliced transcripts per BSJ, it is important to realise that the abundance values of our method are not directly comparable to qPCR results. We thus run CYCLeR on dataset GSE75733 and use the qPCR values reported in (25) for BSJs associated with a unique circular isoform. For this, we average the estimated CYCLeR abundance for two PA1 replicates and correlate it to the average qPCR value per BSJ. This comparison is made with the values reported in (25). Note that CIRIquant is not included in this benchmarking as it cannot handle single-end RNA-seq input data.

#### 2.7.2 Analysis of *D. melanogaster* data

As stated earlier, every tool for circular transcript reconstruction relies on *circRNA* enriched libraries as input to function well. The lack of large, *circRNA* enriched data sets, however, is a major problem in the field. We chose *D. melanogaster* as a model organism for this study due the availability of RNase R treated samples from mature fly head, S2 cell line and early embryo (GSE69212, GSE55872). After creating an index based on the aforementioned samples, CYCLeR quantification is performed on a data set containing 103 *D. melanogaster* samples published by the Lai lab (10). We compare the predictions of CYCLeR to those predicted by representative methods of class I and class III. As representative for class I, we chose the BSJ identification module of CIRCexplorer 2, since this tool has been shown to outperform other tools in class I (2). The representatives of class IIIa produce optimal output when provided with *circRNA* enriched libraries for each sample. Since this dataset does not fit this requirement and as sailfish-cir is the only class III tool that outputs full transcriptome information and simultaneously quantifies linear and circular RNAs, we use sailfish-cir as representative for class III. The observed difference in performance between CYCLeR and sailfish-cir can be attributed to the lack of *de novo* assembly in sailfish-cir. Benchmarking against this tool allows us to highlight the advantage of circRNA transcriptome assembly and its influence on the full-isoform quantification. Note that class IV methods do not work efficiently without *circRNA* enriched data as indicated by simulations.

We normalise BSJ counts from CIRCexplorer 2 as counts-per-billion (CPB) and convert abundance of CYCLeR and sailfish-cir to RPKM. We variance-stabilise the corresponding values using the DESeq2 package and use them to calculate sample adjacencies as Spearman’s rank correlation coefficient. Based on the adjacencies, we calculated sample distances with the topological overlap matrix calculation from the WGCNA package (29).

## 3 RESULTS

In this section we show that CYCLeR outperforms all competing, existing methods based on simulated data. We also highlight the merits of correct isoform information by analysing *D. melanogaster* transcriptome data. Finally, we showcase the difference of output between BSJ-centric tools and CYCLeR and discuss the limitations of qPCR benchmarks.

### 3.1 Reconstruction of circRNA transcripts from simulated data

We had to omit CircAST from our resulting plots, because the script failed to run due to virtual memory issues, even on 400 GB RAM compute nodes of our high-performance computer cluster. We combined the transcripts reconstructed from simulated replicates into one set per tool. For CIRI-full, only the transcripts reconstructed from circRNA enriched libraries are considered.

We separate the simulated transcripts into two reference sets, a low complexity one (that serves as our reference set) and a high complexity one, see supplementary information Figure S2 and Table S2. We show sensitivity and precision plots based on the reference set in Figure 3. Please refer to supplementary information Figure S4 for the sensitivity and precision plot for the corresponding results on the high complexity reference set. For both data sets, CYCLeR clearly outperforms the existing tools both in terms of sensitivity and precision.

**Figure 3:**
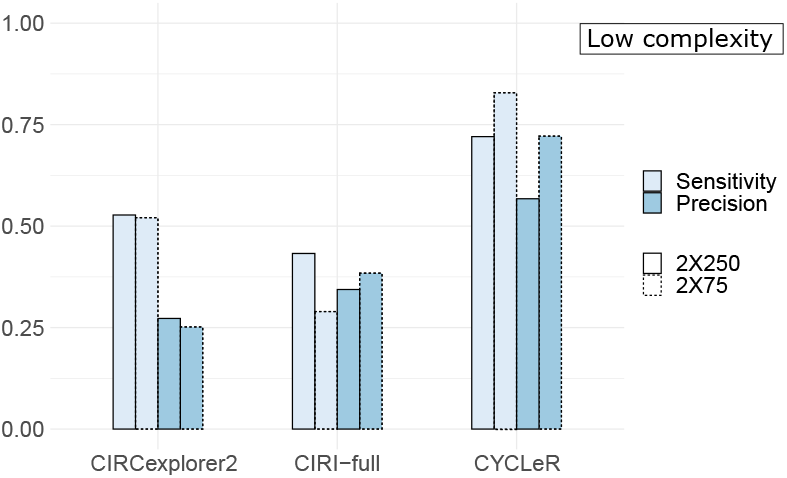
**Comparative benchmarking of** CYCLeR **and selected tools.** Bar plot of sensitivity and precision of CYCLeR and different existing tools based on the simulated reference dataset.CYCLeR outperforms the competitive tools in both sensitivity and precision on both 2X250 and 2X75 sequencing modes.

As can be seen from the benchmarking, CIRI-FULL achieves high precision, but has only limited sensitivity. This is due to the fact that the algorithm outputs the full sequence of *circRNA*s only when there is no break in the coverage of its putative exons. This strategy essentially removes any circle larger than the fragment length of the library. For the 2×75 bp dataset, CIRI-FULL covers mostly cases with single circRNA isoform. For the 2×250 bp dataset, the sensitivity increases, but the precision drops, due to increased complexity of the alternative splicing landscape. CIRCexplorer2 was primarily designed as a tool for detecting alternative splicing events in *circRNA*s and reports as output transcripts corresponding to all potential combinations of splicing events, hence its overall low precision. We observe a minimal difference between the results for the 2×75 bp and the 2×250 bp data sets. In the benchmark with the high complexity 2×250 bp reference set, CIRCexplorer 2 manages to slightly outperform CYCLeR in terms of sensitivity, see supplementary information Figure S4, yet CYCLeR simultaneously significantly outperforms in terms of precision, thereby outperforming CIRCexplorer2 overall.

One advantage of CYCLeR is that it does not have any implicit or explicit limitations in terms of the insert sizes or the read lengths of the RNA-seq library that it can handle. The quantification of genomic features within CYCLeR is also not as reliant on high sequencing depths as the other tools as it solely relies on quantification by junction reads. CYCLeR thereby utilises the entire RNA-seq library for transcript assembly, not only a fraction of around 20% of the reads that happen to span splice sites. This feature of CYCLeR, however, requires a dedicated scaling and normalisation of read counts for exons. We optimise the scaling for reads spanning no more than two exons. In the 2×250 bp dataset, there are naturally more reads spanning multiple exons, thus leading to a decrease in performance of CYCLeR compared to the 2×75 bp set.

Compared to all existing tools, CYCLeR’s strongest advantage is the graph algorithm that is taylor-made for circRNA transcript assembly. A second significant advantage is CYCLeR’s capability to use a combination of BSJ identification tools as input. In its current version, we deliberately chose to exercise extreme caution in reconstruction from low abundance loci. This yields a high precision, but also filters out several low abundance features, i.e. implies a slightly reduced sensitivity, please refer to supplementary information Table S4 for more details in that regard.

### 3.2 CircRNA transcript quantification from simulated data

Table 3 presents the quantification of *circRNA*s from total RNA-seq and RNase R treated RNA-seq simulated data. We designed the simulated dataset in order for the linear transcripts to provide maximum disruption for the correct processing of the circular transcripts. For this benchmark, only class IV tools were considered as only these tools quantify full isoforms, not the BSJ spanning reads only.

**Table 3:**
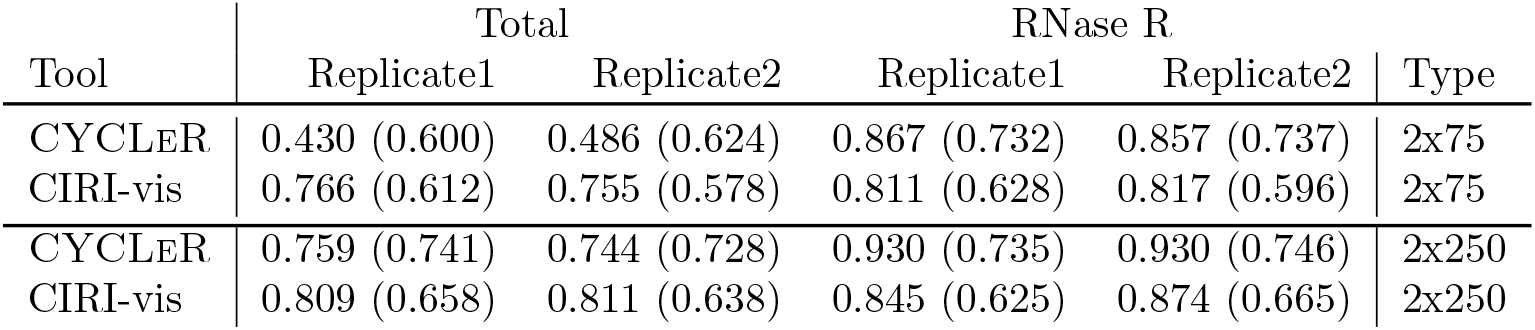
Correlation of predicted versus simulated *circRNA* transcript counts. Correlation of predicted transcript abundances versus simulated. Correlations are based only on the values of correctly identified transcripts. The values are based on correlations for the transcripts of the reference set (in brackets: for transcripts of the the high complexity set)

| Tool | Total |  | RNase R |  | Type |
| --- | --- | --- | --- | --- | --- |
|  | Replicate1 | Replicate2 | Replicate1 | Replicate2 |  |
| CYCLER | 0.430 (0.600) | 0.486 (0.624) | 0.867 (0.732) | 0.857 (0.737) | 2x75 |
| CIRI-vis | 0.766 (0.612) | 0.755 (0.578) | 0.811 (0.628) | 0.817 (0.596) | 2x75 |
| CYCLER | 0.759 (0.741) | 0.744 (0.728) | 0.930 (0.735) | 0.930 (0.746) | 2x250 |
| CIRI-vis | 0.809 (0.658) | 0.811 (0.638) | 0.845 (0.625) | 0.874 (0.665) | 2x250 |

The values in Table 3 are based only on the estimated quantities of the correctly identified transcripts. In this way, we judge the final output of these programs. It is important to note that CYCLeR is the only existing method to simultaneously quantify both known linear and newly assembled circular transcripts. Note that CIRI-vis(3) (the quantification module of the CIRI-suite) is not affected by nascent and linear RNA levels in the same way. Also, CIRI-VIS has lower sensitivity in the total RNA-seq samples, which covers almost exclusively circRNAs simulated without *alternative splicing* events (especially for the 2×75 bp libraries). This leads to CIRI-VIS outperforming in the total RNA-seq samples for low complexity reference set. For the high complexity reference set, the CIRI-VIS output involves more alternative splicing events and – in this case – CYCLeR quantification outperforms all. With multiple overlapping circRNAs, CYCLeR improves its performance, because the impact of the linear and nascent RNA on the reconstruction and henceforth the quantification is lower.

With lower linear and nascent RNA levels in the RNase R treated samples, CYCLeR clearly outperforms CIRI-VIS on all accounts.

### 3.3 Consistency of the assembly

To evaluate the consistency of isoform assembly between samples, we compared the sets of transcripts reconstructed from different samples. The summary of the overlaps is shown in table Table 5. The overlap between biological replicates from the same strain diverges by about 10%. When we compare head RNA-seq libraries between strains, the difference substantially increases. When comparing the reconstructed transcripts between different stages of development or cell lines the overlap is minimal, see supplementary information Figure S9.

**Table 4:** *D. melanogaster* data set: total number of identified transcripts. Summary of the full number of BSJs and transcripts that have been identified by the corresponding tools.

|  | BSJs | Transcripts |
| --- | --- | --- |
| CIRCEXPLORER2 | 12554 | - |
| SAILFISH-CIR | 11117 | 11515 |
| CYCLER | 4371 | 5659 |

**Table 5:** Summary of transcript assembly between different transcriptome samples. Pair-wise set difference and set overlaps between between samples. Column 1 provides information on the 2 samples in the pair-wise comparison of reconstructed transcripts, columns 2 and 4 specify information about the number of different transcripts between samples and column 3 contains the number of overlapping transcripts.

| Sample A & Sample B | A\B | A∩B | B\A |
| --- | --- | --- | --- |
| *D. melanogaster*(Head) RNase R WT19: Rep1 & Rep2 | 311 | 3017 | 298 |
| *D. melanogaster*(Head) RNase R WT28: Rep1 & Rep2 | 199 | 2343 | 196 |
| *D. melanogaster*(Head) RNase R: WT19 & WT28 | 1889 | 1737 | 1001 |
| PA1 cell line: PolyA(-) & PolyA(-)/RNase R | 1003 | 6075 | 761 |

We also compare the reconstruction from different treatments for *circRNA* enrichment. The PA1 cell line has available treatments, only polyA-depletion and a combination of polyA-depletion and RNase R treatment. Naturally, the sample with double depletion substantially increases the number of detected BSJs compared to single depletion – 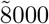 versus 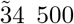. The set of reconstructed transcripts is dependent on the starting set of BSJs. We resolve the conflict by selecting a BSJ set that is derived from the total RNA-seq of the PA1 cell line 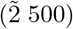, since those are the BSJs that belong to the *circRNA*s quantified later. Based on that set of BSJs, we compare the sets of assembled transcripts. The difference is comparable to the different between biological replicates.

### 3.4 Benchmark with respect to qPCR values

The benchmark from (25) focuses on 13 BSJs. The BSJ locus of CAMSAP1 (Chr9:135881632-135883078), however, has been experimental evidence for two alternative isoforms sharing the same BSJ site, see supplementary information Figure S11A. Based on the output from CYCLeR, we can also conclude that the BSJ locus CORO1C (Chr12:108652271-108654410) yields at least two alternative isoforms, see supplementary information Figure S11B.

Compared to CLEAR, CYCLeR has a lower correlation with qPCR results (0.66 vs 0.75). The difference can be attributed to the fact that CYCLeR quantification is more affected by GC and length biases. The results of the abundance estimations can be seen in supplementary information Figure S10. CIRIquant is not included in this benchmark as it cannot handle single-end RNA-seq data.

### 3.5 Analysis of *D. melanogaster* data

In addition to investigating the merits of CYCLeR when dealing with simulated data, where the isoforms and their respective abundances are known up-front, we are also keen to explore CYCLeR and other tools when analysing real transcriptome data. For this, we investigate RNA-seq data from *D. melanogaster*, see the section above for more details.

For CIRCexplorer 2, we normalise the counts as CPM, whereas we normalise those from sailfish-cir and CYCLeR as RPKM. All counts are then variance stabilised using VST from the DESeq2 package (30) It is important to note the number of BSJs that are included in this analysis, see the numbers in Table 4. CIRCexplorer 2 includes all the BSJs identified in the analysis, whereas sailfish-cir filters out the BSJs that are not part of the linear annotation. CYCLeR uses the BSJs that correspond to the RNase R treated dataset.

Figure 4 (A) and (B) show the UMAP dimensional scaling (31) of all 103 samples from the dataset of the Lai lab. There is no indication of overall loss of information due to the decrease of the BSJ set. The procedure is also repeated for sailfish-cir and shown in Figure S7. Within the dataset, the subset with most time points corresponds to different stages during embryo development. The embryo development samples were extracted and the UMAP scaling was repeated, see also supplementary information Figure S8. It is apparent that the data is heavily affected by multiple batch effects. We annotated part of the batches based on SRA ascension numbers.

**Figure 4:**
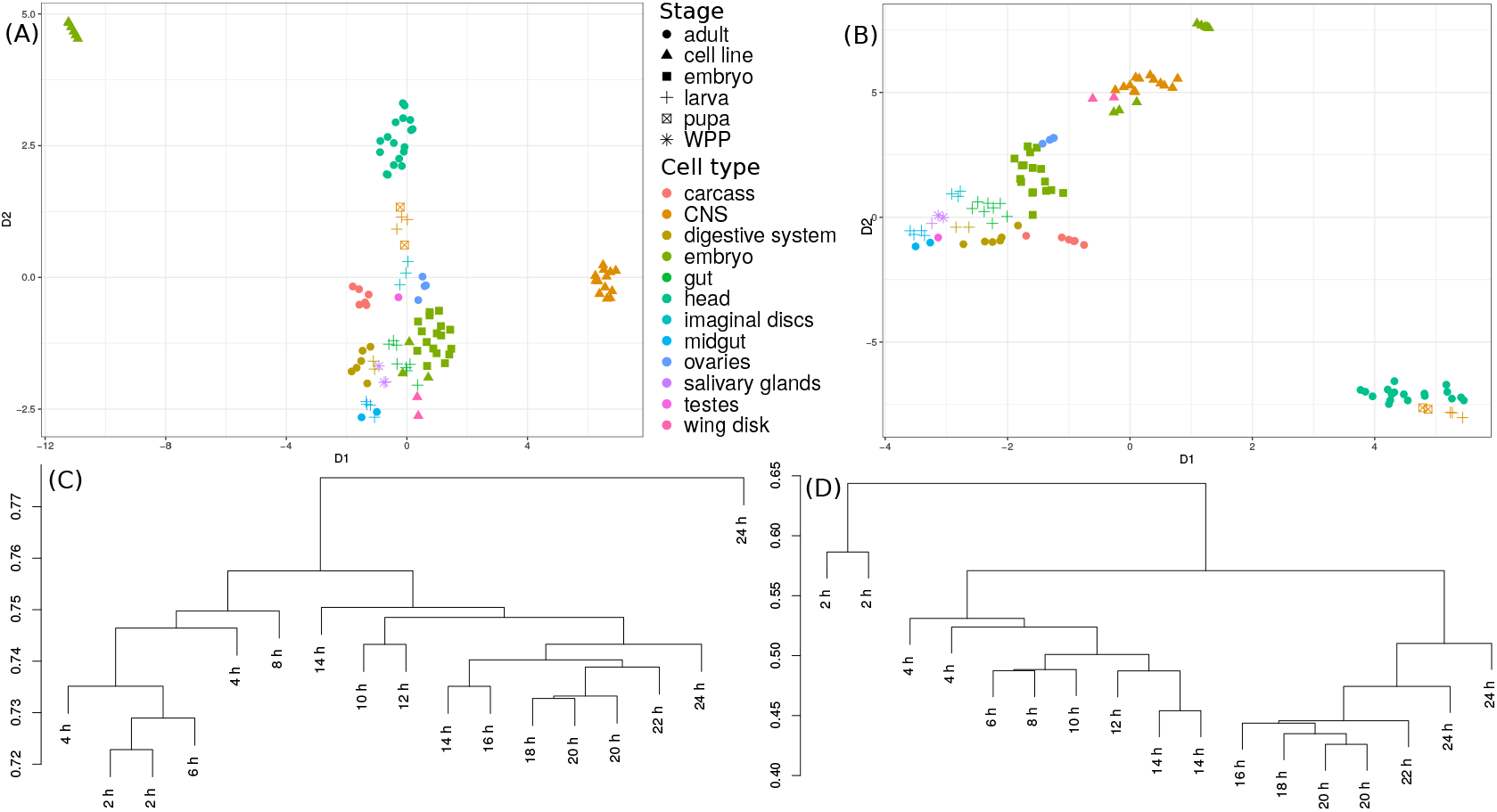
**Comparison of** CIRCexplorer**2 and** CYCLeR **for the *D. melanogaster* transcriptome sets.** The comparison is made based exclusively on circRNA abundance estimations. (A) and (B) show the UMAP dimensional scaling of the abundances inferred by CIRCEXPLORER2’s BSJ detection module and CYCLeR for all 103 samples of the Lai dataset. (C) (CIRCexplorer2) and (D) (CYCLeR) show a dendrogram of the subset of data corresponding to embryo stages which are based on between sample distance calculations. The scale of the dendrograms represents the samples’ distances.

There are two noteworthy differences to observe. Using CYCLeR for quantification, it is possible to identify a gradient in the data that reproduces the known developmental stages. In addition, the quantification by CYCLeR makes the outliers in the data easily distinguishable. The reason for this advantage of CYCLeR derives from its variance stability between replicates, see supplementary information Figure S6. This difference can also be clearly seen within the dendrogram representing the sample distances based on circRNA transcript counts, see Figure 4 (C) and (D). The clustering by CYCLeR is in complete agreement with the results from (7) depicting two segregated stages of fly embryo development, namely pre-mbl expression and post-mbl expression. The corresponding MBL protein is known to affect *circRNA* expression by binding intronic sequences. The most notable difference in the *circRNA* profile is circ-mbl (exon2) becoming the most expressed *circRNA*. Overall, the distances between replicate samples inferred by CYCLeR are significantly more biologically meaningful, emphasising that the correct assembly of full-sequence isoforms is key for the correct clustering of samples.

## 4 DISCUSSION

We here present CYCLeR, the first computational method for identifying the full sequence identify of new *circRNA*s and their abundances while simultaneously co-estimating the abundances of known linear splicing isoforms. These linear isoforms can be specified as optional input by the user.

The common Bioinformatics approach towards identifying circRNAs is primarily based on tracing the levels of BSJ spanning reads across different samples. Some of the existing tools supplement this BSJ information by ratios of BSJ/FSJ spanning reads (25; 2). This forces the user to implement circRNA-specific analytical pipelines (2). Even these analytical pipelines, however, cannot distinguish between alternatively spliced circRNAs mapping to the same BSJ. While the BSJ/FSJ ratio is very consistent with qPCR results for circRNAs comprising two exons (25), we still require methods that are able to identify the full sequence identity of all *circRNA*s and their alternative splice variants in order to be able to apply standard differential expression analyses or clustering pipelines. CIRI-full provides full isoform sequence, but the algorithm cannot reconstruct and quantify the isoforms efficiently from non-enriched data. This forces the user to perform statistical analyses of the circRNA based primarily on the BSJ-spanning reads, even when the full isoform data is available. Also, the circular and linear transcripts need to be quantified separately and subsequently combined in a co-expression network(4). CYCLeR simultaneously quantifies linear and circular transcripts from the same sample in an integrated manner, thereby giving the user the opportunity to proceed with standard downstream pipelines for quantitative analysis. This is essential for enabling co-expression analyses of linear and circular RNAs, since wrong abundances can severely impair optimal clustering.

CYCLeR outperforms existing tools for *circRNA* identification and quantification on all accounts on simulated data. Our reference set provides a benchmark similar to existing *circRNA* studies and allows us to gain a better understanding of tool-specific biases caused by the challenging mapping of chimeric fragments. Our high complexity data set was devised to represent the full complexity of real data, where the low abundances of transcripts and the overlap between multiple circular and linear isoforms can render the identification of the full isoform profile a near impossible task. When dealing with complex data, the true value of the tool becomes less centered around the recall and more focused on precision. The full set of circRNA isoforms is impossible to reconstruct due to natural limitation imposed by the depth of the RNA-seq library. The algorithm of CYCLeR ensures that the full sequence identity of the most abundant isoforms is correctly reconstructed. This advantage of the assembly allows for reliable quantification of the predominant isoforms. The quantification of *circRNA*s in CYCLeR is enhanced by the fact that assembly and abundance estimation are performed in separate steps. This makes CYCLeR very robust, as erroneously assembled transcripts are estimated with significantly lower abundance than the correctly assembled ones. Existing tools that depend on long library insert size typically fail to detect *circRNA*s that fall below the detection limit of the libraries (22). In contrast to this, CYCLeR is independent of library insert size, allowing for short insert sizes and retaining the ability to identify both short and long *circRNA*s.

The experimental workload required by analyses performed with CYCLeR is substantially lower than for alternative methods. We show in a study of *D. melanogaster* transcriptomes that samples with RNase R treatment for a few key time points are sufficient to produce biologically meaningful results. We also showcase the merits of proper *circRNA* isoform detection for correct *circRNA* quantification. Finally, we find that CYCLeR improves samples clustering and facilitates outlier sample detection. This is an important feature that will play a key role in the technological transition towards single cell experiments.

## References

1. Xin, R., Gao, Y., Gao, Y., Wang, R., Kadash-Edmondson, K.E., Liu, B., Wang, Y., Lin, L. and Xing, Y.(2021) isoCirc catalogs full-length circular RNA isoforms in human transcriptomes. Nat. Commun., 12, 1–11.

2. Zhang, J., Chen, S., Yang, J. and Zhao, F. (2020) Accurate quantification of circular RNAs identifies extensive circular isoform switching events. Nat. Commun., 11,.

3. Zheng, Y., Zhao, F. and Zhao, F. (2020) Visualization of circular RNAs and their internal splicing events from transcriptomic data. Bioinformatics, 36, 2934–2935.

4. Ji, P., Wu, W., Chen, S., Zheng, Y., Zhou, L., Zhang, J., Cheng, H., Yan, J., Zhang, S., Yang, P., et al.(2019) Expanded Expression Landscape and Prioritization of Circular RNAs in Mammals. Cell Rep., 26, 3444–3460.e5.

5. Pandey, P.R., Rout, P.K., Das, A., Gorospe, M. and Panda, A.C.(2019) RPAD (RNase R treatment, polyadenylation, and poly(A)+ RNA depletion) method to isolate highly pure circular RNA. Methods, 155, 41–48.

6. Cape, B., Swain, A., Nicolis, S., Hacker, A., Walter, M., and Koopman, P. (1993) Circular Transcripts of the Testis-Determining Gene Sry in Adult Mouse Testis. Cell, 73, 1019–1030.

7. Ashwal-fluss, R., Meyer, M., Pamudurti, N.R., Ivanov, A., Bartok, O., Hanan, M., Evantal, N., Memczak, S., Rajewsky, N., and Kadener, S. (2014) Article circRNA Biogenesis Competes with Pre-mRNA Splicing. MOLCEL, 56(1), 55–66.

8. Memczak, S., Jens, M., Elefsinioti, A., Torti, F., Krueger, J., Rybak, A., Maier, L., Mackowiak, S.D., Gregersen, L.H., Munschauer, M., Loewer, A., Ziebold, U., Landthaler, M., Kocks, C., Noble, F., and Rajewsky, N. (2013) Circular RNAs are a large class of animal RNAs with regulatory potency. Nature, 495(7441), 333–338.

9. Salzman, J., Gawad, C., Wang, P.L., Lacayo, N., and Brown, P.O. (2012) Circular RNAs Are the Predominant Transcript Isoform from Hundreds of Human Genes in Diverse Cell Types. PLoS ONE, 7(2), e30733.

10. Westholm, J.O., Miura, P., Graveley, B.R., Lai, E.C., Westholm, J.O., Miura, P., Olson, S., Shenker, S., Joseph, B., and Sanfilippo, P. (2014) Genome-wide Analysis of Drosophila Circular RNAs Reveals Their Structural and Sequence Properties and Age-Dependent Neural Accumulation. CellReports, 9(5), 1966–1980.

11. Zhang, X., Wang, H.b., Zhang, Y., Lu, X., Chen, L.l., and Yang, L. (2014) Complementary Sequence-Mediated Exon Circularization. Cell, 159(1), 134–147.

12. Li, Z., Huang, C., Bao, C., Chen, L., Lin, M., Wang, X., Zhong, G., Yu, B., Hu, W., Dai, L., Zhu, P., Chang, Z., Wu, Q., Zhao, Y., Jia, Y., Xu, P., Liu, H., and Shan, G. (2016) Exon-intron circular RNAs regulate transcription in the nucleus. Nat. Struct. Mol. Biol., 22(3), 256–264.

13. Zhou, C., Molinie, B., Daneshvar, K., Pondick, J.V., Wang, J., Wittenberghe, O.V., Xing, Y., Giallourakis, C.C., and Mullen, A.C. (2017) Identification and characterization of m6A circular RNA epitranscriptomes. bioRxiv, 1–45.

14. Yang, Y., Fan, X., Mao, M., Song, X., Wu, P., Zhang, Y., and Jin, Y. (2017) Extensive translation of circular RNAs driven by N6-methyladenosine. Nat. Publ. Gr., 27(5), 626–641.

15. Trapnell, C., Williams, B.A., Pertea, G., Mortazavi, A., Kwan, G., van Baren, M.J., Salzberg, S.L., Wold, B.J., and Pachter, L. (2010) Transcript assembly and quantification by RNA-Seq reveals unannotated transcripts and isoform switching during cell differentiation. Nat. Biotechnol., 28(5), 511–5.

16. Pertea, M., Pertea, G.M., Antonescu, C.M., Chang, T.C., Mendell, J.T., and Salzberg, S.L. (2015) StringTie enables improved reconstruction of a transcriptome from RNA-seq reads. Nat. Biotechnol., 33(3), 290–295.

17. Szabo, L. and Salzman, J. (2016) Detecting circular RNAs: bioinformatic and experimental challenges. Nat. Rev. Genet., 17(11), 679–692.

18. Jeck, W.R., Sorrentino, J.A., Wang, K.A.I., Slevin, M.K., Burd, C.E., Liu, J., Marzluff, W.F., and Sharpless, N.E. (2013) Circular RNAs are abundant, conserved, and associated with ALU repeats. RNA, 141–157.

19. Zhang, X.o., Dong, R., Zhang, Y., Zhang, J.l., Luo, Z., Zhang, J., Chen, L.l., and Yang, L. (2016) Diverse alternative back-splicing and alternative splicing landscape of circular RNAs. Genome Res., 1277–1287.

20. Gao, Y., Wang, J., Zheng, Y., Zhang, J., Chen, S., and Zhao, F. (2016) Comprehensive identification of internal structure and alternative splicing events in circular RNAs. Nat. Commun., 7(May), 1–13.

21. Metge, F., Czaja-hasse, L.F., Reinhardt, R., and Dieterich, C. (2017) FUCHS — towards full circular RNA characterization using RNAseq. PeerJ, 1–14.

22. Zheng, Y., Ji, P., Chen, S., Hou, L., and Zhao, F. (2019) Reconstruction of full-length circular RNAs enables isoform-level quantification. Genome Med., 11(1), 1–20.

23. Li, M., Xie, X., Zhou, J., Sheng, M., Yin, X., Ko, E.a., Zhou, T., and Gu, W. (2017) Quantifying circular RNA expression from RNA-seq data using model-based framework. Bioinformatics, 1–9.

24. Wu, J., Li, Y., Wang, C., Cui, Y., Xu, T., Wang, C., Wang, X., Sha, J., Jiang, B., Wang, K., Hu, Z., Guo, X., and Song, X. (2019) CircAST: Full-length Assembly and Quantification of Alternatively Spliced Isoforms in Circular RNAs. Genomics, Proteomics and Bioinformatics, 17(5), 522–534.

25. Ma, X.K., Wang, M.R., Liu, C.X., Dong, R., Carmichael, G.G., Chen, L.L., and Yang, L. (2019) CIRCexplorer3: A CLEAR Pipeline for Direct Comparison of Circular and Linear RNA Expression. Genomics, Proteomics Bioinforma., 17(5), 511–521.

26. Li, H. (2013) Aligning sequence reads, clone sequences and assembly contigs with BWA-MEM. arXiv, 00(00), 1–3.

27. Dobin, A., Davis, C.A., Schlesinger, F., Drenkow, J., Zaleski, C., Jha, S., Batut, P., Chaisson, M., and Gingeras, T.R. (2013) STAR: Ultrafast universal RNA-seq aligner. Bioinformatics, 29(1), 15–21.

28. Bray, N.L., Pimentel, H., Melsted, P., and Pachter, L. (2016) Near-optimal probabilistic RNA-seq quantification. Nat. Biotechnol., 34(5), 525–527.

29. Langfelder, P. and Horvath, S. (2008) WGCNA: an R package for weighted correlation network analysis. BMC Bioinformatics, 9(1), 559.

30. Love, M.I., Huber, W., and Anders, S. (2014) Moderated estimation of fold change and dispersion for RNA-seq data with DESeq2. Genome Biol., 15(12), 1–21.

31. McInnes, L., Healy, J., and Melville, J. (2018) UMAP: Uniform Manifold Approximation and Projection for Dimension Reduction. arXiv, 1–51.

32. Kim, D., Pertea, G., Trapnell, C., Pimentel, H., Kelley, R., and Salzberg, S.L. (2013) TopHat2: accurate alignment of transcriptomes in the presence of insertions, deletions and gene fusions. Genome Biol., 14(4), 1–13.

33. Kim, D., Langmead, B., and Salzberg, S.L. (2015) HISAT: a fast spliced aligner with low memory requirements. Nat. Methods, 12(4), 357–360.

34. Anders, S., Reyes, a., and Huber, W. (2012) Detecting diferential usage of exons from RNA-seq data. Genome Res, 22(10), 2008–2017.

35. Goldstein, L.D., Cao, Y., Pau, G., Lawrence, M., Wu, T.D., Seshagiri, S., and Gentleman, R. (2016) Prediction and quantification of splice events from RNA-seq data. PLoS ONE, 11(5), 1–18.

